# The effects of clay minerals on bacterial community composition during arthropod decay

**DOI:** 10.1101/2024.02.19.580992

**Authors:** Nora Corthésy, Farid Saleh, Camille Thomas, Jonathan B. Antcliffe, Allison C. Daley

## Abstract

Fossilization, or the transition of an organism from the biosphere to the geosphere, is a complex mechanism involving numerous biological and geological variables. Bacteria are one of the most significant biotic players to decompose organic matter in natural environments, early on during fossilization. However, bacterial processes are difficult to characterize as many different abiotic conditions can influence bacterial efficiency in degrading tissues. One potentially important variable is the composition and nature of the sediment on which a carcass is deposited after death. We experimentally examined this by decaying the marine shrimp *Palaemon varians* underwater on three different clay sediments. Samples were then analyzed using 16S ribosomal RNA sequencing to identify the bacterial communities associated with each clay system. Results show that samples decaying on the surface of kaolinite have a lower bacterial diversity than those decaying on the surface of bentonite and montmorillonite, which could explain the limited decay of carcasses deposited on this clay. However, this is not the only role played by kaolinite, as a greater proportion of gram-negative over gram-positive bacteria is observed in this system. Gram-positive bacteria are generally thought to be more efficient at recycling complex polysaccharides such as those forming the body walls of arthropods. This is the first experimental evidence of sediments shaping an entire bacterial community. Such interaction between sediments and bacteria might have contributed to arthropods’ exquisite preservation and prevalence in kaolinite-rich Lagerstätten of the Cambrian Explosion.

## INTRODUCTION

Microorganisms, are ubiquitous and play crucial roles in numerous processes, including animal metabolism, biogeochemical cycles, and the degradation of organic matter (Azam et al., 1994; Fava, 2015; Madsen, 2011). For a long time, little was known about the animal microbiome, which is primarily responsible for digestion and immunity, but it has recently gained ground in research, largely due to high-throughput sequencing (McAdam et al., 2014). Studying bacteria is essential because, in many systems, such as the human body, bacterial cell quantities exceed those of host cells (Sender et al., 2016). Given the widespread presence of bacteria, they are recognized as one of the primary biotic factors governing the post-mortem decay of biological tissues, with a preference for recycling soft anatomical structures (e.g., skin, internal organs, non-mineralized cuticles) over hard and resistant tissues (e.g., bones, horns, claws, mineralized shells) (Nudds & Selden, 2008).

The bacterial processes involved in decay and fossilization are poorly understood as they include many complex interactions between bacteria, the nature of the decaying organic matter, in addition to the substrate in the depositional environment. For example, experiments on the decomposition of brine shrimps have revealed that the gut remains well-preserved, despite hosting numerous microorganisms (Butler et al., 2015). This discovery suggests that the gut microbiota might be crucial for preserving internal organs and therefore could play a role in the preservation of soft tissues in the fossil record (Butler et al., 2015). Indeed, exceptionally preserved animal fossils from several Cambrian Burgess Shale-type Lagerstätten sometimes show the presence of high-relief three-dimension gut structures preserved in iron oxide or phosphatic minerals, while the rest of the labile anatomy is preserved as flattened carbon films (Butterfield, 2002; Lerosey-Aubril et al., 2012; Vannier et al., 2014). Many mineralization processes, in which soft tissues are replicated in minerals like pyrite or phosphate crystals, resistant over geological time scales, are mediated or accelerated by bacterial activity (Saleh et al., 2020). Therefore, bacteria are not only responsible for the decay of soft tissues but can also stabilize labile anatomical features, via their metabolic pathways under certain conditions. One approach to understanding the role of bacteria in degradation, given their limited preservation potential in the rock record (Saleh et al., 2020; Vinther, 2016), is through controlled decay experimentation.

Many previous decay experiments have investigated the impact of numerous abiotic factors on the rate and sequence of organic matter degradation. These factors include oxygen (Butler et al., 2015; Hancy & Antcliffe, 2020; Murdock et al., 2014; Sansom, 2016; Sansom et al., 2010), pH (Clements et al., 2017, 2022), and associated sediment composition (Martin et al., 2004; Naimark et al., 2016; Sagemann et al., 1999; Wilson & Butterfield, 2014). Experiments such as these have provided an outstanding framework of the sequences of anatomical character loss and how that varies under different conditions. It is now time to ask why these variations are occurring and to describe the processes leading to their observed differences. Decay experiments involving sediments have shown that kaolinite, a clay mineral often associated with Cambrian Burgess Shale-type preservation (Anderson et al., 2018, 2021), preserves organisms better than other substrates (Naimark et al., 2016; Wilson & Butterfield, 2014). By inoculating a single species of bacteria into a kaolinite suspension, it has been shown that this clay can limit bacterial growth probably due to its high aluminum concentration, which would damage bacterial membranes in the presence of oxygen (McMahon et al., 2016). However, generalization should be carefully made as recent microbiome analysis of decaying crayfish, placed separately in aerobic and anaerobic conditions, revealed the presence of diverse bacterial communities around the decaying carcasses (Mähler et al., 2023). Therefore, we know that kaolinite can affect bacteria (McMahon et al., 2016). However, we do not know how kaolinite influences the composition of the bacterial communities within and surrounding a decaying carcass (Mähler et al., 2023), and the process by which kaolinite aids exceptional fossil preservation is yet to be fully deciphered (Naimark et al., 2016; Wilson & Butterfield, 2014).

In this study, bacterial communities associated with the marine shrimps *Palaemon varians* decaying on three different clay substrates were examined using 16S ribosomal RNA (rRNA) sequencing. The results show that the bacterial communities present in the shrimp and the sediments on which it is decaying depend on the carcass and its surrounding substrate. Kaolinite impacts soft-tissue degradation by promoting the proliferation of bacteria that are less efficient at recycling polysaccharide-rich body walls, such as those of arthropods. The observation of contrasting bacterial communities in association with different clays begins to provide a process-based understanding as to why certain animals in Early Paleozoic ecosystems are so exquisitely preserved in rock matrices containing kaolinite.

## MATERIAL AND METHODS

### Sample preparation

To investigate the interplay of clay minerals and bacterial communities during animal decomposition, the decay of the marine shrimp *Palaemon varians* was studied. Seventeen marine shrimps were euthanized using clove oil (C_7_H_12_CIN_3_O_2_) at the aquarium lab facility of the Institute of Earth Sciences at the University of Lausanne. Clove oil was chosen to avoid mechanically damaging the animals and to induce their rapid death. After three minutes, all animals were successfully euthanized, and they were repeatedly rinsed with deionized water to remove clove oil residue. Experimental boxes (5x3x2cm) were prepared in the meantime and contained 5g dry weight of clay powder and 35g of artificial seawater (ASW), prepared to 1.024 psu with reverse osmosis deionized water and Aquarium Systems Reef Crystals. The three clays used in the experimental setup were kaolinite, bentonite, and montmorillonite (Pusch, 2015; Saikia et al., 2003; Uddin, 2008). The carcasses were then individually placed on the surface of the clay in the decay experiment boxes, which were then closed with lids to limit oxygen supply and prevent excessive evaporation. Control samples of 5g clay powder with 35g of ASW without any carcass were also prepared. Controls were prepared for each clay, resulting in three additional samples. All samples were kept at room temperature and in the dark to avoid bias in microbial growth (Sansom, 2014). Shrimps were left to decay for two months. Specimen were then imaged using a SC50 5-megapixel color camera (Olympus Life Science Solutions) with Olympus Stream Basic software (version 2.2; Olympus Soft Imaging Solutions). Six shrimp samples (two decaying on each clay mineral), in addition to the three controls, were prepared for DNA extractions. The investigated shrimps were intentionally not sterilized as the experiment aims to explore the influence of different clay minerals on both the bacteria present within the shrimp (e.g., in its guts) and environmental bacteria (on its cuticle). After two months, the remaining pieces of the shrimp carcass and the surrounding sediment were subsampled in sterile 1.5ml tubes. For the controls, only sediments were put into tubes. Tubes were then stored in the freezer at -18°C to limit the decay process and were kept frozen until DNA extractions. We acknowledge that this experimental setup does not replicate natural environments, as a single type of clays does not form the entire seafloor, and other environmental factors such as oxygen concentrations and the presence of scavengers are not accounted for. However, it is important to note that the primary focus of decay experiments is not to mimic natural conditions but rather to investigate how specific variables influence decay patterns. In this case, the variable under investigation is sediment mineralogy, and its impact on shrimp decay is assessed through the analysis of bacterial communities. Given the impossibility of replicating natural environments, especially those from half a billion years ago, we chose not to inoculate our experiment with any randomly occurring microbial community found in nature. This decision avoids increasing the number of variables investigated and prevents potential distortion from our main objective, which is to examine the influence of shrimp bacteria on decay in the presence of different clays.

### DNA extractions and 16S sequencing

DNA extractions of the nine samples were conducted at the Department of Earth Sciences of University of Geneva. The DNeasy PowerSoil Pro Kit (Qiagen, Germantown, MD, USA), one of the most common kits for the extraction of DNA from sedimentary environments, was used following the manufacturer’s protocol. For each tube, any water remaining in the samples was removed by sterile pipetting. For the control samples, 500mg of the clay was placed individually into a PowerBead Pro Tube. For samples containing shrimp carcasses, pieces of the carcass were harvested with surrounding clays and placed in the same types of tubes until reaching 500mg. The masses of each remaining carcass were not measured. The standard protocol was then followed. The DNA was quantified by using Invitrogen Qubit dsDNA HS assay kit (Life Technologies, #Q32851, Grand Island, NY, USA) and was subsequently stored at -18°C. An extraction blank, without sample material, was realized during sample manipulation and DNA extractions to assess potential microbial contaminations. The 16S rRNA gene amplification and sequencing were conducted by Macrogen Inc. (Seoul, Rep. of Korea). The V3-V4 region of the 16S rRNA gene was amplified by primers 341F (5’-CCTACGGGNGGCWGCAG-3’) and 805R (5’-CCCCGYCAATTCMTTTRAGT-3’; Herlemann et al., 2011) as these primers are often used for shrimp gut microbial identification and recommended for good coverage of bacterial diversity (García-López et al., 2020; Holt et al., 2021). The V3-V4 16S rRNA amplicons were sequenced using an Illumina MiSeq (paired-reads run 2 × 300 bp, San Diego, CA, USA).

### Microbiome analysis

Microbiome analyses were performed using *dada2* (version 1.25.2; Callahan et al., 2016) R package. Taxonomic affiliation was done against the SILVA database version nr99_v138.1, (Quast et al., 2013). Data were then analyzed using *phyloseq* (version 1.36.0; (McMurdie & Holmes, 2013)), *vegan* (version 2.6-4) and *ggplot2* (version 3.4.0; (Wickham, 2016)) on RStudio. The generated barplots facilitate in-depth investigation between samples with shrimps and controls, as well as a more detailed exploration of the differences in bacterial communities between the different minerals. The bacterial diversity of each sample was assessed with the Shannon-Weaver index at the amplicon sequence variant (ASV) level. Similarity and dissimilarity of the bacterial communities between samples were also assessed at the ASV level with a non-metric multidimensional scaling (NMDS) plot based on a Bray-Curtis distance metric after their normalization using a Hellinger transformation. Then, a permutational analysis of variance (ANOVA) was done using Bray-Curtis dissimilarity to test if bacterial communities were significantly different between the three types of clay, and between kaolinite and the other two combined clays (bentonite and montmorillonite). The results of the ANOVA are influenced by the small sample sizes in this study, a limitation we acknowledge. Significant differences are only deemed present when clear disparities are evident in the plotted data. However, to address the challenge posed by the numerous variables in the system (i.e., the large number of bacterial taxa) and the limited number of specimens, we elected to categorize the bacteria into two distinct groups based on their cell wall composition: gram-positive and gram-negative bacteria. The two categories were defined at the class level (Tab. 1), only considering the classes whose relative abundances exceed 1%. For the classes that may have gram-positive or gram-negative cell walls (e.g., Bacilli, Clostridia), the categorization was done at the order level (Tab. 1). ANOVA was then used, once more, to test whether the relative abundances of gram-positive and gram-negative bacteria were significantly different between the three individual clays, and between kaolinite and the two other combined clays.

**Table 1.** Bacterial phyla, classes, and orders identified in the samples, and their type of cell walls (gram-positive/gram-negative).

| Phyla | Classes and *orders* | Gram |
| --- | --- | --- |
| Actinobacteriota | Actinobacteria | Positive |
| Bacteroidota | Bacteroidia | Negative |
| Desulfobacterota | Desulfobulbia | Negative |
|  | Desulfovibrionia | Negative |
| Firmicutes | Bacilli |  |
|  | *Lactobacillales* | Positive |
|  | Clostridia |  |
|  | *Clostridiaceae* | Positive |
| Fusobacteriota | Fusobacteriia | Negative |
| Planctomycetota | Phycisphaerae | Negative |
| Proteobacteria | Alphaproteobacteria | Negative |
|  | Gammaproteobacteria | Negative |

When the interaction between the type of clay and the type of cell wall was significant, contrast analyses were performed with *emmeans* (version 1.8.2) R package to assess in more detail whether the proportions of gram-positive and gram-negative bacteria were influenced by the different clays. Raw datasets and statistical analyses can be found in the *Supplementary Materials* and downloaded on the following link (https://doi.org/10.17605/osf.io/K6DHG).

## RESULTS AND DISUCUSSION

### Bacterial diversity is individual sample and clay dependent

The phylum-scale data show that Firmicutes and Desulfobacterota are less abundant in the presence of kaolinite than in the presence of the two other clays (Fig. 1A). The proportions of Bacteroidota and Planctomycetota (Wiegand et al., 2018) are higher in the presence of kaolinite in comparison to the bentonite and montmorillonite experiments (Fig. 1A). When comparing the three control groups only (i.e., without a decaying shrimps), differences in bacterial communities are also observed. In particular, the control sample containing kaolinite shows a lower Firmicutes presence compared to the controls with bentonite/montmorillonite (Fig. 1A). All these observations suggest that clays influence bacterial community composition, with kaolinite playing a different role compared to bentonite and montmorillonite.

**Figure 1.**
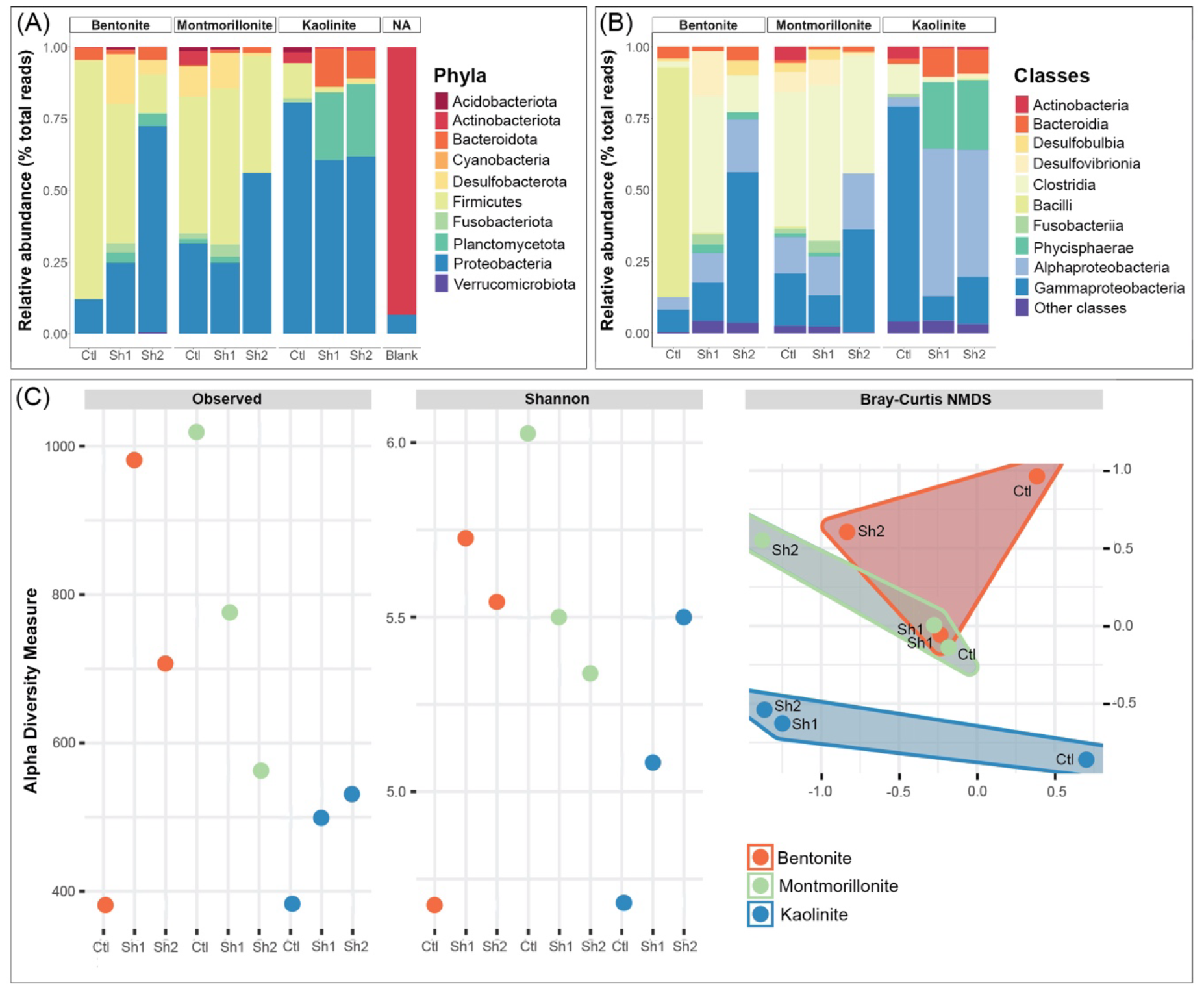
(A) Relative bacterial abundances at the phylum level in the presence of different clay minerals. Each clay barplot includes one control (Ctl) consisting of sediment without shrimp and two marine shrimps (Sh1 and Sh2) decaying for two months. The blank represents contaminations during DNA extraction and manipulation. (B) Relative bacterial abundances at the class level in the presence of different clay minerals. (C) Species level raw alpha diversity and Shannon diversity index in the presence of different clay minerals in addition to NMDS analyses showing an overlap between bentonite and montmorillonite with both plotting separately from kaolinite.

Comparisons between the shrimps decaying on bentonite reveal remarkable variation between the two individual samples, even though they were placed on the same substrate (Fig. 1A). A similar observation is made for the two shrimps placed on montmorillonite (Fig. 1A). This highlights that shrimps decaying under identical experimental conditions can exhibit variations in their bacterial communities, which implies that bacterial composition is influenced by the decaying individual and its unique microbiome. Nonetheless, the impact of the individual sample on bacterial diversity appears relatively minimal when kaolinite is present, as the two shrimps decaying on kaolinite share a comparable bacterial composition (Fig. 1A). Although the possibility that the two shrimps placed on kaolinite had a similar original bacterial composition cannot be completely ruled out, these results also suggest that kaolinite may have a more pronounced impact on the bacterial community composition than the other clays, leading to a homogenization of the bacterial diversity in kaolinite shrimp samples in our study (Fig. 1A).

Differences between kaolinite and the other clays are also evident when examining bacterial classes (Fig. 1B). Notably, the kaolinite samples display an almost absence of Clostridia members from the Firmicutes phylum (Fig. 1B). Within the Proteobacteria phylum, the kaolinite samples are marked by the prevalence of Alphaproteobacteria over Gammaproteobacteria when shrimps are present in the experiment (Fig. 1B). Additionally, the kaolinite samples exhibit an increased presence of Phycisphaerae and Bacteroidia members compared to bentonite and montmorillonite. In contrast, bentonite and montmorillonite display a higher relative abundance of Desulfobulbia and Desulfovibrionia within the Desulfobacterota phylum (Fig. 1B). All the samples differ substantially from the blank, which consists mainly of Actinobacteriota (Fig. 1A) and confirms that the obtained data do not result from external contaminations. The observed phyla are consistent with those observed in other decay experiments (Mähler et al., 2023) and with bacteria recovered in arthropod taxa (García-López et al., 2020; Holt et al., 2021).

These findings suggest that none of the investigated clays totally inhibits bacterial activity as all samples show diverse bacterial communities (Fig. 1A, B). However, the reduction of the activity of specific bacterial species by certain clays, as suggested by McMahon et al. (2016), cannot be ruled out. When looking at species-level (ASV) diversity metrics, sequencing read numbers are the lowest for the kaolinite experiments with less than 550 reads for both shrimp-containing samples (Fig. 1C). Similarly, the diversity, as measured by the Shannon-Weaver index is on average lower for the kaolinite shrimp experiments (5.0) when compared to the bentonite and montmorillonite experiments (5.5) (Fig. 1C). The NMDS also shows a marked difference between the kaolinite bacterial communities and the bentonite and montmorillonite communities (Fig. 1C). The ANOVA testing the contrast in bacterial communities between the three clays did not show significant differences (*F*_*1,8*_ = 0.885, *p* = 0.577), and this is likely due to the overlap between bentonite and montmorillonite values (Fig. 1C). However, when combining bentonite and montmorillonite in one group and comparing it to kaolinite, bacterial communities are shown to be significantly different (*F*_*1,8*_ = 2.013, *p* = 0.0304), which supports that samples with kaolinite have meaningfully different communities (Fig. 1C). Note that differences between the three different clays, and possibly between montmorillonite and bentonite, might be more evident if a bigger sample size was considered.

An intriguing observation in our data is that the controls do not cluster together in the NMDS plot, but rather remain proximate to their respective clay-mineral groups (Fig. 1D). This suggests that the clay minerals not only influence the bacteria initially introduced with the shrimp but also any bacteria that could have developed over the two months in the controls. Put differently, the influence of minerals on bacteria could be discerned when comparing the controls to one another. If different clays did not exert differential impacts on bacterial community compositions, all controls would exhibit the same bacterial composition, which is not the case (Fig. 1A, B). However, not all the observations of this experiment could have been made without the samples that included shrimps. For instance, there is also a clear difference between the controls and the shrimp samples for both bentonite and kaolinite (Fig. 1B, C), indicating that shrimps also influence the diversity in the system. This difference is less apparent between montmorillonite samples with shrimps and the montmorillonite control (Fig. 1B). Additionally, the montmorillonite controls show a higher number of reads than the samples with shrimps (Fig. 1C). It is unclear why this might be happening in the montmorillonite control, and more experiments involving montmorillonite need to be conducted in the future.

At this stage, due to its chemical composition, kaolinite may have favored the growth of certain bacteria while inhibiting others, resulting in different bacterial communities than in the presence of bentonite and montmorillonite (Fig. 1D). As a result, the proportions and abundances of both phyla and classes are remarkably different between kaolinite and the other clays (Fig. 1), and it is important to explore the implications of these variances in a decay context.

### Bacteria associated with kaolinite are characteristic of early decay stages

The differences in bacterial compositions between kaolinite, bentonite, and montmorillonite are related to different stages of decay, as bacterial communities change with time (i.e., bacterial succession) after the death of an organism. In the literature, a high abundance of Desulfobacterota (Waite et al., 2020), gammaproteobacterial Enterobacterales and Clostridia are often associated with advanced carcass decay, unlike Bacteroidia and Alphaproteobacteria that are often active during early stages of decomposition (Adserias-Garriga, Quijada, et al., 2017). As highlighted in the previous section, Clostridia and Desulfobacterota are almost absent from the kaolinite samples (Fig. 1A, B). Moreover, Enterobacterales are largely dominant in the gammaproteobacterial sequences of bentonite (66% and 91% of Gammaproteobacteria in Sh1 and Sh2 respectively) and montmorillonite (53% and 43% of Gammaproteobacteria in Sh1 and Sh2 respectively) experiments (see the external link in *Material and Methods* for detailed composition). Therefore, the prevalence of Alphaproteobacteria and Bacteroidia in kaolinite samples (Fig. 1B), as opposed to the dominance of Clostridia (Fig. 1B), Desulfobacterota (Fig. 1A), and gammaproteobacterial Enterobacterales in bentonite and montmorillonite, suggests that the decomposition of shrimp organic matter is at an earlier stage in the kaolinite samples than in the other clays.

Bacterial occurrences are consistent with morphological observations of the decaying shrimps as carcasses are better preserved when they are left to decay on kaolinite than when they decay on bentonite and montmorillonite (Fig. 2). In the presence of bentonite and montmorillonite, all that remains after two months are severely decayed fragments (Fig. 2C, D), while the anatomical parts of shrimps on kaolinite are still discernable (Fig. 2A, B). By limiting the establishment of bacterial communities that are associated with advanced decay, such as the ones observed in bentonite and montmorillonite samples (Adserias-Garriga, Hernández, et al., 2017; Adserias-Garriga, Quijada, et al., 2017; DeBruyn & Hauther, 2017), kaolinite better preserves arthropod carcasses for prolonged periods (Fig. 2A, B). However, the process by which kaolinite drives a specific bacterial community, which is not efficient in recycling shrimp carcasses, remains unknown. A separation of the observed bacteria between gram-negative and gram-positive (Tab. 1) can provide a better understanding of the interactions between clays and different bacterial cell walls and allows the investigation of bacterial metabolism in greater depth by limiting the number of variables.

**Figure 2.**
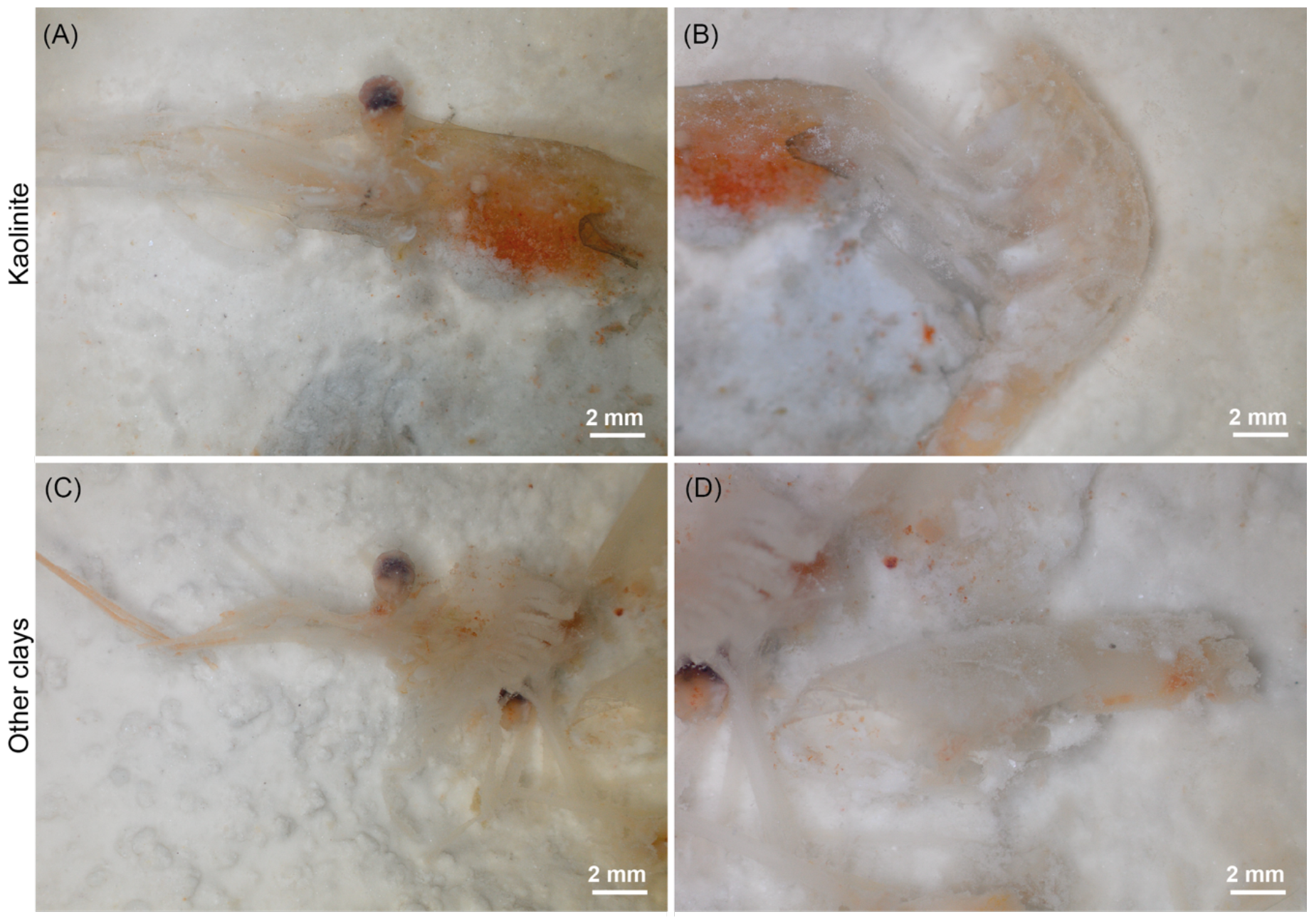
The decay stage of marine shrimps after two months following death when they are deposited on (A, B) kaolinite and on (C, D) the two other clays (bentonite or montmorillonite).

### Kaolinite favors gram-negative over gram-positive bacteria

The relative abundances of gram-positive and gram-negative bacteria are similar for bentonite and montmorillonite samples as each group of bacteria represents approximately 50% of the community (Fig. 3A; p_Bentonite[Neg-Pos]_ = 0.875, t-ratio_B[Neg-Pos]_ = 0.161, p_Montmorillonite[Neg-Pos]_ = 0.927, t-ratio_M[Neg-Pos]_ = 0.093, Tab. S1-S2). However, the proportions of gram-positive and gram-negative bacteria for the samples with kaolinite are significantly different with a very high proportion of gram-negative and a very low proportion of gram-positive bacteria (Fig. 3A; p_Kaolinite[Neg-Pos]_ = 0.0003, t-ratio_K[Neg-Pos]_ = 4.966, Table S1-S2). When comparing the proportions of gram-positive bacteria between kaolinite samples and all samples from the other two clays, the proportions are significantly lower for kaolinite than for the other clays (Fig. 3B; p_Positive[Kaolinite-Others]_ = 0.0089, t-ratio_Pos[Kaolinite-Others]_ = -3.034, Tab. S3-S4). In contrast, the proportions of gram-negative bacteria for kaolinite are significantly higher than for the other two clays (Fig. 3B; p_Negative[Kaolinite-Others]_ = 0.0096, t-ratio_Neg[Kaolinite-Others]_ = 2.999, Tab. S3-S4). The difference in the proportions of gram-negative and gram-positive bacteria in the presence of kaolinite can be explained by the fact that gram-positive bacteria are more sensitive to high aluminum concentrations and thus to kaolinite than are gram-negative bacteria (Jou & Malek, 2016; Piña & Cervantes, 1996).

**Figure 3.**
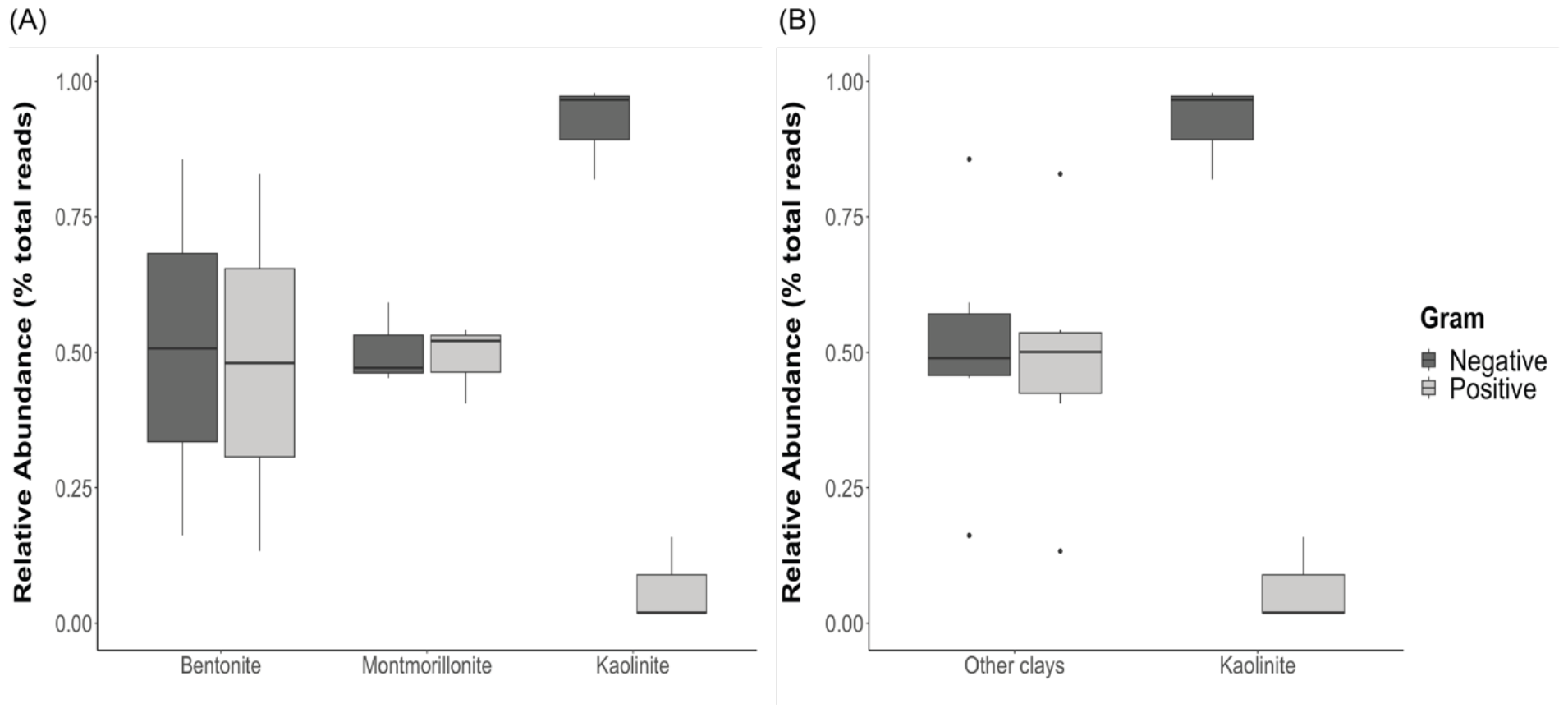
(A) Relative abundances of gram-negative and gram-positive bacteria in the presence of the three different clay minerals. (B) Relative abundances of gram-negative and gram-positive bacteria in the presence of bentonite and montmorillonite combined (other clays) and kaolinite. Each boxplot includes the proportion of gram-negative or gram-positive bacteria, defined at the class level, of the two shrimp samples and the control corresponding to the tested clay.

Both gram-negative and gram-positive bacteria have been identified in association with decaying organic material (Zhou et al., 2021). Community fingerprinting from soils of different organic matter composition and vegetation stages showed that gram-positive to gram-negative bacteria ratio was high in soils with complex carbon chains (Fanin et al., 2019). In soils, it was suggested that gram-positive bacteria mainly rely on recalcitrant complex carbon compounds as an energy source, whereas gram-negative bacteria use simpler carbon compounds (Kramer & Gleixner, 2008). The cuticle of shrimps, composed of chitin, consists of a polysaccharide with complex carbon chains (Baas et al., 1995; Das et al., 2016) and would typically serve as an abundant carbon source for gram-positive bacteria. However, since kaolinite is likely to favor the growth of gram-negative bacteria (Fig. 3), it would result in a reduced recycling of chitin and an increase in the preservation potential of shrimp exoskeletal elements in the presence of kaolinite (Fig. 2A, B) providing novel insights into the process of their exceptional fossilization.

The positive impact of kaolinite on the preservation of polysaccharidic body walls may extend beyond arthropods to other cuticularized animal groups such as annelids and priapulids. Arthropods, priapulids, and annelids are prominent constituents of modern ecosystems, as they were around half a billion years ago during the Cambrian and the Ordovician (Daley et al., 2018; Nanglu et al., 2020; Saleh et al., 2021, 2022). Many Early Paleozoic biotas with these animal groups are preserved in sediments containing kaolinite or other reactive minerals (Anderson et al., 2018, 2021; Saleh et al., 2019). Although the favorable role of kaolinite can be attributed to its high aluminum concentration (Amonette et al., 2003; Guida et al., 1991; Imlay et al., 1988), it is expected that different types of kaolinite, with varying elemental compositions, may have varying effects on bacterial community compositions. Furthermore, the beneficial impact on exceptional preservation may not be exclusive to kaolinite. It is possible that other clays and minerals could fulfill a similar role if they possess specific physico-chemical properties, such as aluminum enrichment. It will be interesting in the future to expand analyses to try new clay minerals, and other animal groups than arthropods. However, at this stage, it is possible to suggest that the preservation of some ecdysozoa and lophotrocozoa in the fossil record might be facilitated by kaolinite, through its role in favoring gram-negative over gram-positive bacteria during organic matter decay.

## CONCLUSION

Bacterial community compositions associated with decaying shrimps deposited on the surface of three different clays were investigated. Results show that bacterial composition is influenced by both the decaying organism and the surrounding clay (Fig. 1A, B). None of the investigated clays completely inhibits bacterial activity (Fig. 1A, B), but the kaolinite system shows a lower bacterial diversity in comparison to the systems with bentonite and montmorillonite (Fig. 1C). Importantly, a significant majority of bacteria associated with kaolinite are gram-negative (Fig. 3). The preferential proliferation of gram-negative over gram-positive bacteria in the presence of kaolinite may imply that complex carbon chains cannot be efficiently recycled when this clay is present. This phenomenon can explain why the body walls of organisms like shrimps, which are composed of polysaccharides, exhibit better preservation in kaolinite compared to when other clays are present (Fig. 2). These results start to explain the pattern observed in the fossil record whereby exceptionally well-preserved arthropods from the Cambrian are often found associated with kaolinite-rich sediments, by showing that the preservation was likely linked to the effect of this clay mineral on bacterial communities in such a way as to slow the decay of the carcass.

## Acknowledgments

The authors thank fruitful discussions with all members of the Anom Lab at the University of Lausanne, Switzerland, and with numerous attendees of the PalAss Annual Meeting in Cambridge, UK.

## Funding

NC and FS are funded by an SNF Ambizione Grant (PZ00P2_209102). JBA is supported by an SNF Sinergia Grant (198691) awarded to ACD. Lab and experimental expenses were supported by the University of Lausanne, the University of Geneva, and an SNF project grant (205321_179084) awarded to ACD.

## Conflict of interests

The authors declare no competing interests.

## Data availability

All data necessary to replicate this work are available in the main text, in the *Supplementary Materials*, and on the following link (https://doi.org/10.17605/osf.io/K6DHG).

## Author contributions

NC, JBA, and ACD designed the research. NC and CT conducted the experiments. NC and FS interpreted and discussed the results with all co-authors. NC made the figures and wrote the initial version of the text with the help of all co-authors.

## Notes

### Competing Interest Statement

The authors have declared no competing interest.

