## Supplementary Material for "The effects of clay minerals on bacterial community composition during arthropod decay"

**Raw data and processed of the microbiome analyses can be found on the following link** (https://doi.org/10.17605/osf.io/K6DHG).

### Detailed statistical analyses

**Table S1.** ANOVA comparing bacterial relative abundances according to clay minerals and cell walls (gram-positive/gram-negative), of the three clays.

| | *DF* | *Sum square* | *Mean square* | *F-value* | *p-value* |
|---|---|---|---|---|---|
| Clay | 2 | $4 \cdot 10^{-5}$ | $2 \cdot 10^{-5}$ | $5 \cdot 10^{-4}$ | 0.999 |
| Gram | 1 | 0.404 | 0.404 | 9.083 | 0.011 |
| Clay*Gram | 2 | 0.695 | 0.348 | 7.804 | 0.007 |
| Residuals | 12 | 0.534 | 0.045 | | |

**Table S2.** Contrast analyses to assess whether the proportions of gram-positive/gram-negative bacteria are influenced by the three different clays.

| | *Estimate* | *Standard Error* | *t-ratio* | *p-value* |
|---|---|---|---|---|
| **Clay = Bentonite** | | | | |
| Negative – Positive | 0.028 | 0.172 | 0.161 | 0.875 |
| **Clay = Kaolinite** | | | | |
| Negative – Positive | 0.856 | 0.172 | 4.966 | 0.0003 |
| **Clay = Montmorillonite** | | | | |
| Negative – Positive | 0.016 | 0.172 | 0.093 | 0.927 |
| **Gram = Negative** | | | | |
| Bentonite – Kaolinite | -0.413 | 0.172 | -2.396 | 0.080 |
| Bentonite – Montmorillonite | 0.003 | 0.172 | 0.019 | 0.999 |
| Kaolinite – Montmorillonite | 0.416 | 0.172 | 2.415 | 0.078 |
| **Gram = Positive** | | | | |
| Bentonite – Kaolinite | 0.415 | 0.172 | 2.408 | 0.079 |
| Bentonite – Montmorillonite | -0.008 | 0.172 | -0.049 | 0.999 |
| Kaolinite – Montmorillonite | -0.423 | 0.172 | -2.458 | 0.072 |

**Table S3.** ANOVA comparing bacterial relative abundances according to clay minerals and cell walls (gram-positive/gram-negative), when comparing kaolinite with bentonite and montmorillonite combined.

| | *DF* | *Sum square* | *Mean square* | *F-value* | *p-value* |
|---|---|---|---|---|---|
| Clay | 1 | $2 \cdot 10^{-5}$ | $2 \cdot 10^{-5}$ | $6 \cdot 10^{-4}$ | 0.981 |
| Gram | 1 | 0.404 | 0.404 | 10.594 | 0.006 |
| Clay*Gram | 1 | 0.695 | 0.695 | 18.203 | 0.001 |
| Residuals | 14 | 0.534 | 0.038 | | |

**Table S4.** Contrast analyses to assess whether the proportions of gram-positive/gram-negative bacteria are influenced by the different clays, when comparing kaolinite with bentonite and montmorillonite combined.

| | *Estimate* | *Standard Error* | *t-ratio* | *p-value* |
|---|---|---|---|---|
| **Clay = Kaolinite** | | | | |
| Negative – Positive | 0.856 | 0.160 | 5.363 | 0.0001 |
| **Clay = Other clays** | | | | |
| Negative – Positive | 0.022 | 0.113 | 0.194 | 0.849 |
| **Gram = Negative** | | | | |
| Kaolinite – Other Clays | 0.414 | 0.138 | 2.999 | 0.0096 |
| **Gram = Positive** | | | | |
| Kaolinite – Other Clays | -0.419 | 0.138 | -3.034 | 0.0089 |
